# Habitat edges affect tree diversity more than biomass regeneration in a reforested wet neotropical timber plantation

**DOI:** 10.1101/2022.12.06.515700

**Authors:** Nicholas Medina, Emma Capetz, Andrea Johnson, Agustín Mendoza, Max Villalobos

## Abstract

About half of all forests are tropical and secondary, making tropical forest regeneration integral to future forests. Tree stand biomass and taxonomic richness can recover in a few decades, but relative abundances may lag indefinitely. Since most forests are within a km of a habitat edge, edge effects likely affect community composition regeneration. However, most studies assess how degraded edges affect intact forests, leaving it unclear whether higher-quality edges could facilitate regeneration of nearby degraded forests. Notably, higher quality edges near intact forests could promote processes like dispersal and wood biomass accumulation that effectively accelerate succession, leading to better performance of shade-tolerant taxa compared to pioneer taxa in the early stages of forested plantation regeneration. This study addressed how wet tropical forested plantation regeneration was affected by distance to adjacent intact forest edge. It was hypothesized that old timber plantations facilitate regeneration by increasing available shade, favoring the presence and biomass of later-successional taxa, ultimately changing community composition overall. A wet neotropical timber plantation reforested after 20 years and adjacent to primary forest was censused for trees along a 300 m edge distance gradient, and analysis matched identified taxa to broad dispersal mode and wood density traits using relevant literature.

As distance from primary forest edge increased, stem and wood density tended to increase significantly, with ~10% variation explained, while biomass and canopy light surprisingly tended to stay the same. Stand tree richness also tended to increase significantly, but diversity decreased steeply and non-linearly, explained in part by wood density, and taxonomic composition varied notably. Finally, tree taxa associated with both early and late successional stages decreased significantly, as well as genus Ficus, but biomass by dispersal mode did not tend to change. Overall this study supports that stand composition is less resilient and more subject to edge effects than biomass and richness, suggesting that global forests will likely be distinctly new assemblages in the future, with timber and biodiversity trade-offs occurring based on local and regional management activity.

## Introduction

Forest and landscape restoration is a key international conservation and climate change adaptation strategy (*De Pinto et al., 2020*). While tropical forests specifically store most land biomass (*Pan et al., 2013*), most are now secondary (*FAO, 2020*) and functionally degraded (*Hubau et al., 2020*). This likely amplifies amid increasing climate stressors (*Anderegg et al., 2022*) and collective management issues, including insufficient policy support (*Chazdon, 2018*) via skewed priorities (*Pyron and Mooers, 2022*). Overall, secondary forests can regenerate relatively quickly compared to old-growth forests, in some ways. Neotropical secondary forests can grow quickly enough to accumulate biomass >10x faster than old-growth forests (*Finegan, 1996; Poorter et al., 2016*), and substantially offset carbon emissions, with estimates ranging from ~10% from Amazonian deforestation (*Smith et al., 2020*) to a decade of fossil fuel emissions from all of Latin American and the Caribbean (*Chazdon et al., 2016*). They also recover taxonomic richness and biomass quite quickly, as well as tree height, averaging 80%of old-growth forest levels in just 20 years, especially in wetter regions (*Rozendaal et al., 2019*). The recovery of species richness alone is beneficial in that it also tends to correlate with the recovery of some ecosystem services (*Guariguata and Ostertag, 2001; Stanturf et al.,2014*) like biomass and carbon storage (*Liu et al., 2018*). However, recovery of secondary neotropical forests diverge widely in taxonomic composition (*Norden et al., 2015*), potentially taking over a century to recover (*Poorter et al., 2021*) with added variability (*Atkinson et al., 2022*). Despite being one of the slowest ecosystem properties to recover, restoring recovering community composition is often important for preserving rare taxa (*Carlo and Morales, 2016*), which can be keystone to locally-adapted food webs, including birds (*Maas et al., 2016*). Overall, understanding secondary forest regeneration is key for global biodiversity conservation at the global scale.

Modern secondary forest regeneration is often affected by edge effects, or changes near an ecosystem boundary, especially in heterogeneously-managed landscapes (*Melo et al., 2013; Perfecto et al., 2009*). About 70% of all forests have been estimated to be within a km of their edge (*Haddad et al., 2015*), resulting in tropical forests having been estimated to lose ~22% aboveground carbon along heavily-managed edges >100m, in part due to shortening trees and lighter leaf mass (*Ordway and Asner, 2020*). Effects of degraded edges have historically favored tree taxa that tend to be pioneering, or be faster-growing, have lower wood densities, and be less shade-tolerant (*Tabarelli et al., 2008*). Accordingly, secondary forest wood density may also tend to decrease near degraded habitat edges. Additionally, the dominance of pioneering taxa could lead to arrested succession (*Tymen et al., 2016*), which further delays the recruitment of shade-tolerant taxa, as well as associated increases in aboveground biomass, in part due to their higher survival with increasing shade, and tendency toward relatively higher wood densities. However, arrested succession could be prevented, and even typical succession accelerated, if intact forests that were adjacent to regenerating plantations functioned as contributing reservoirs of seeds from later-successional tree taxa. In this case, having a habitat edge near a relatively rich intact forest, rather than near a relatively degraded open area, would instead be an asset to regenerating plantation forest patches, rather than a detriment. Overall, edge effects ultimately have the potential to shape community composition, in part based on the local dispersal kernel patterns of existing taxa (*Wandrag et al., 2017*), which while complicated (*Muller-Landau and Hardesty, 2009*), can also correlate with simpler plant traits like height and seed mass (*Tamme et al., 2014; Thomson et al., 2011*).

Additionally, properties of the tropical forest carbon stores and tree community depend heavily on land-use history and previous management (*Omeja et al., 2012; Pyles et al., 2022*). Overall, legacy effects from management like intensive timber planting could potentially delay or result in non-linear changes between successional stages (*Albrich et al., 2020; Gough et al., 2021*), which would make future management more uncertain, unlike more sustainable thinning practices and agroforestry (*Lefland et al., 2018*). In some cases, the leftover timber stands could provide additional shade, which would not be available in a clear-cut harvested timber plantation, and this could help more shade-tolerant taxa establish and recruit better. Overall, edge effects near degraded habitat edges are often studied as negative, but conceptual influences from meta-population (*Levins, 1979*) and meta-community (*Leibold et al., 2004; Warren et al., 2015*) theories do highlight dispersal as a key process for offsetting extinction debt and thus maintaining biodiversity, thereby potentially making particular edge effects, specifically from adjacent intact forests, actually beneficial for regeneration plantation forest parcels. Taken together, these ideas suggest that edge effects specifically from adjacent intact primary forests, potentially via shade-tolerant dispersal and performance, may indeed facilitate plantation forest regeneration. Such a beneficial edge effect may ultimately help explain relatively high biodiversity, compared to adjacent stands, found in agroforests (*Oliveira-Neto et al., 2017*), forests affected by logging (*Clark and Covey, 2012; Edwards et al., 2014*), and timber plantations (*Pryde et al., 2015*).

Influenced by restoration ecology for forest management, this study reports tree regeneration patterns of an abandoned wet neotropical timber plantation, focused on highlighting edge effects from an adjacent primary forest on overall stand properties, tree community composition, and limited functional trait recovery. We hypothesized that remaining timber trees would mediate forest regrowth by maintaining shade, which would suppress less shade-tolerant pioneer tree taxa, and in turn facilitate more shade-tolerant tree taxa, ultimately allowing for more tree diversity within the regenerating plantation forest in parts that were closer to the richer edge of the intact unmanaged forest. Accordingly, we predicted more specifically that further from the intact primary forest edge, canopy light availability would increase, while stand biomass and diversity would decrease.

## Methods

### Study site

This study was done in a neotropical lowland rainforest on the Osa Peninsula at the Greg Gund Conservation Center (approx. 8.3778176, −83.2935426) near the Piro Biological Station run by Osa Conservation. (See *Taylor et al. (2015)* for a broader ecosystem description of the region.) The study site was a regrowing 20 ha timber plantation of *Bombacompsis quinata* abandoned in ~1990 after the vulnerably dry-adapted species from the Guanacaste region (*Hulshof and Powers, 2020*) grew poorly in very wet conditions. This focal secondary forest area was roughly triangular, surrounded by primary forest on the two S and NW sides (Fig 1) and a wide service road on the third NE border, with primary forest beyond it. This census was done in 2013 during the rainy season months between June and August.

**Figure 1:**
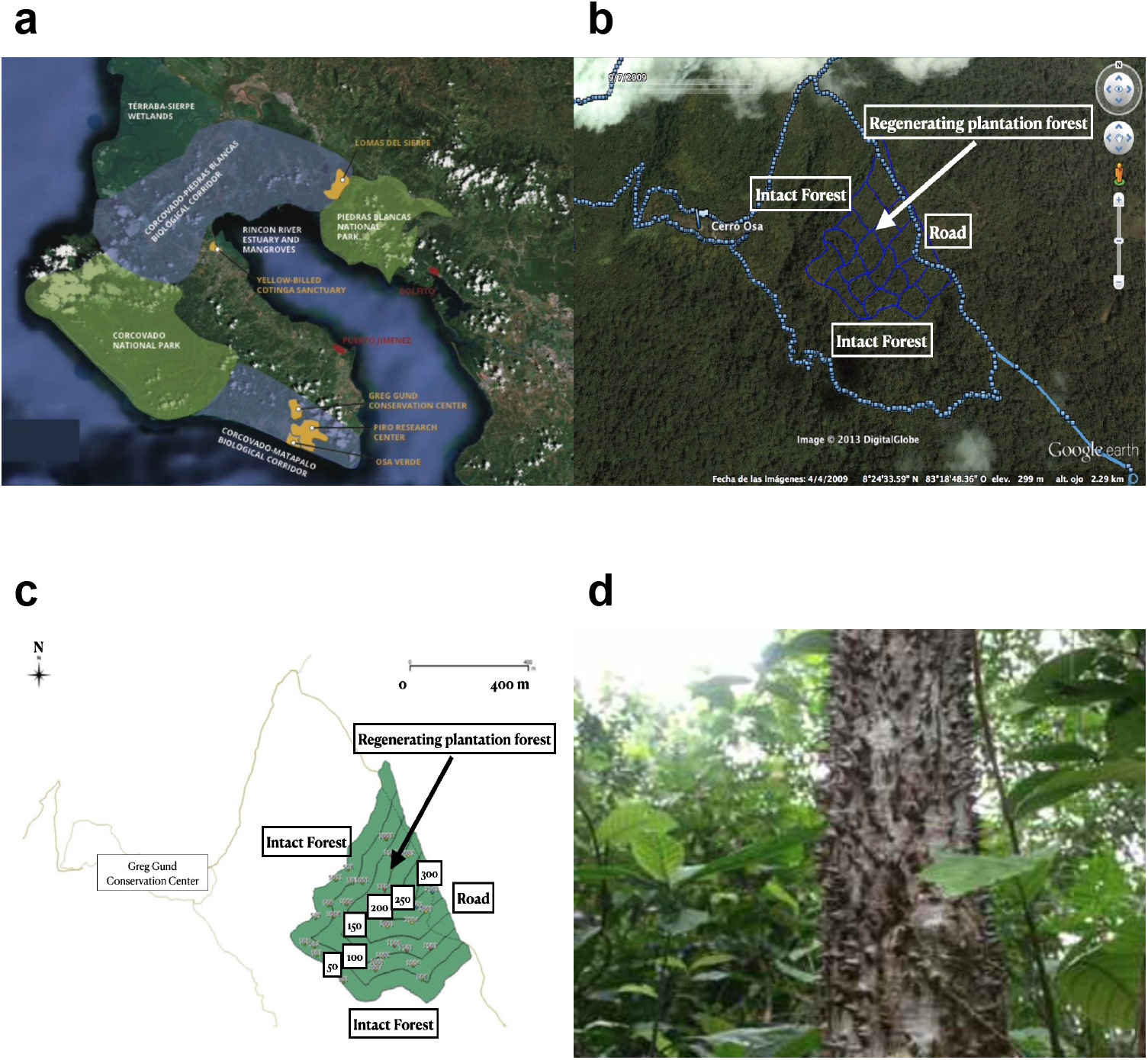
Map of (a) conservation areas and (b) study site (c) showing plot design with 50 m distance strata bins, and (d) focal plantation timber taxon Bombacopsis quinata on Osa Peninsula, Costa Rica. Map image data © 2022 Osa Conservation, © 2009 Google. Map GIS data and photo credit: Max Villalobos, Nicholas Medina.

### Census design

Edge effects were studied by dividing the secondary forest area into six discrete 50 m strata spanning 0 - 300 m away from the primary forest edge on the S and NW sides, going inward to the E (Fig 1), using available GIS software (*ArcGIS 10*, esri.com, and *QGIS 2*, qgis.org). Note that in line with hypotheses about facilitated regeneration, this study most often referred to the “edge” of the regenerating plantation forest as the part closest to the unmanaged intact forest, where trees may be dispersing from, rather than the open service road, which is usually the focus of other studies evaluating effects of forest fragmentation and degradation, rather than restoration. Each stratum was randomly filled with a number of 21 × 21 m square census plots oriented N that was proportional to its area–specifically with 11, eight, five, three, two, and one plot(s), respectively, as distance away from primary forest increased. Given the small size of the last 300 m distance stratum (Fig 1c), one census plot was sufficiently representative. All data were later centered at the stratum level, so resulting trends were not weighted by distance bands with more plots in them. The total area of the 30 census plots equaled ~1 ha or 5% of the total secondary forest stand area, which is comparable to similar studies (*Onyekwelu and Olabiwonnu, 2016*).

### Plot measurements

Light reaching the forest floor was measured at the center of each plot at chest height using a densiometer (*Forestry Suppliers, Inc*.), as an average of four readings taken facing each cardinal direction. The slope of the forest floor was measured using a rangefinder (*Bushnell, Forestry Suppliers, Inc*) to measure the distance the diagonal between two plot corners and triangulate the observation angle.

The diameter of all stems >10 cm wide were recorded in each census plot, totaling over 1,243 trees. In cases where a tree split into 2 or more stems below breast height, each stem was measured separately; in cases where a stem split only above breast height, it was measured as a single stem. Tree height was recorded by measuring distances to both the crown and the stem at chest height (*~2.7 m*) using a rangefinder (*Bushnell, Forestry Suppliers, Inc*.) and triangulating the missing side length.

Taxa were identified by local experts, and trait information was gathered from the literature. Final traits used only included successional stage (early, late) and main dispersal mode (wind, animal, water), which were ultimately matched to species using only the specific dataset made available by *Werden et al. (2020)*. Otherwise wood tissue carbon was matched to genera with the GLOWCAD database (*Doraisami et al., 2022*), given high taxonomic alignment of this trait above the species level (*Chave et al., 2006*).

Aboveground biomass (AGB) was calculated at the individual stem level using the allometric equation by *Chave et al. (2015)* and later centered at the plot level. The equation specifically for wet tropical forests was used, which has performed slightly better compared to both pantropical and previous other models by (*Alvarez et al., 2012; Chave et al., 2005; Ngomanda et al., 2014*). The equation version with height was used since height was accessibly measurable and has shown improved local accuracy (*Domke et al., 2012*). Wood density (or specific gravity) values when available also improve biomass estimates even more than height (*Van Breugel et al., 2011*). The equation used was:

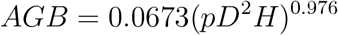

with diameter *D* in cm, height *H* in m, and wood density or specific gravity *p* in *g cm*^-3^. DBHs and heights were measured in the field and wood densities were extracted from available literature. Values were updated using the pipeline developed in the *BIOMASS* 2.1.8 R software package by *Réjou-Méchain et al. (2017)*, which notably replaced unknown wood density values with plot-level averages instead during stem biomass calculations.

### Statistical analyses

For all response variables, data were centered using medians at the plot level, followed by the discrete distance stratum level where applicable, which was most cases and indicated on figures. These medians were then subjected to linear regression with distance to intact forest edge as the only independent variable, which was binned discretely according to census design. All regressions were run through assumption checks of residual normality with Shapiro-Wilk tests and equal variance with Levene tests, using base R version 4.2.1 (2022-06-23) and *rstatix* 0.7.0 package functions (*R Core Team, 2022*).

Non-linear regressions were run using the function *poly()* in the base R package *stats* 4.2.1, which was only ultimately recorded for stem density and richness variables, based on significance and explained variance over linear models. All trees censused were included in each analysis, including unidentified taxa, which were only grouped together for taxonomic analyses and whose exclusion did not affect observed patterns shown in results. Community analysis and ordination was done as a PERMANOVA with the *adonis()* function from the *vegan* 2.6.4 R package (*Dixon, 2003*). Data and code were organized with R packages *here (Müller, 2020), bookdown (Xie, 2022*), and *grateful (Rodríguez-Sánchez et al., 2022*), and internal pipeline *oir (Medina, 2022a*), and are stored at github.com/nmedina17/osa (*Medina, 2022b*) and https://doi.org/10.5281/zenodo.7406478 (*Medina, 2022c*).

## Results

### Biomass

Aboveground biomass did not tend to change with distance to primary forest edge (Fig 2a), despite significant trends with underlying related variables (Fig 2). Median plot wood density tended to increase marginally significantly (p=0.105) with edge distance by 0.00019 per m explaining *~10* % of variance among median distance strata values (Fig 2b), while tissue carbon did not change significantly with distance to forest edge (Fig 2c). Plot stem density also tended to increase significantly (p=0.03) with edge distance but more strongly and non-linearly, by ~17.39 ± 3.4 per m (Fig 2d). At this stage in forest regeneration, canopy light availability did not tend to change with distance from forest edge (Fig 2e). Notably, tree diversity decreased marginally significantly (p=<0.0001) by ~ 1 per g plot wood density, yet only explaining *~4* % of variance (Fig 2f). Additional variables tested, including maximum height and diameter, tended to stay the same across the edge gradient (*data not shown*).

~~~
## |                                                                                         |
~~~

**Figure 2:**
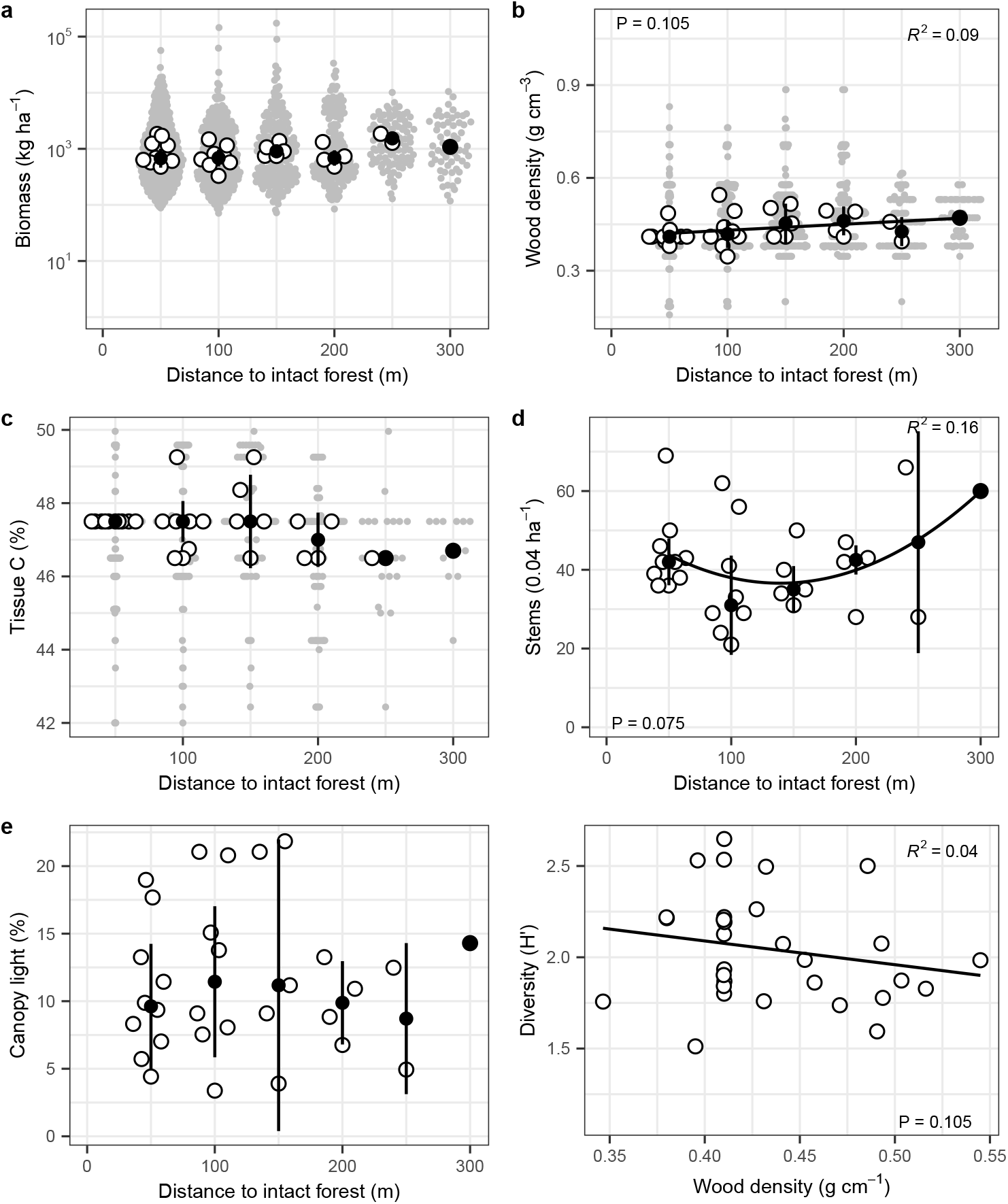
Plot stand properties, namely (a) biomass, (b) wood density, (c) tissue carbon, (d) stem density, (e) canopy light availability, all across distance to primary forest edge gradient, and (f) diversity against wood density. Grey dots show individual tree values, white dots with border show plot medians, black dots show plot-level data binned into discrete distance strata medians ± 1 absolute deviation, and black lines show linear or non-linear regression through strata medians.

### Composition

Overall diversity changed significantly with distance to primary forest edge (Fig 3). Diversity significantly (p=0.003) decreased non-linearly by ~0.5 ± 0.029 per m as distance to primary forest edge increased, which explained *73*% of variance among distance strata medians (Fig 3a) – while in contrast taxonomic richness increased slightly (p=0.067) and linearly by 0.0063 ± 0.0025 per m and had *16*% variance explained among distance strata (Fig 3b). Community composition and beta diversity also changed significantly (p=0.01) with 11.38% variance among distance strata explained by distance to primary forest edge, and the first two principal components explaining 47% and 23% totaling 70% of variance among calculated plot distances (Fig 3d). Key abundant taxa *Vochysia* and *Ficus* showed different responses – *Vochysia* nearly tended to decrease with distance to primary forest edge (Fig 3e), while *Ficus* decreased marginally significantly (p=0.088) and linearly by −62.62 ± 25.05 per m with *~20*% variance among distance strata explained (Fig 3f). Other taxa did not change significantly with distance to intact forest (*data not shown*).

~~~
## |                                                                                         |
~~~

**Figure 3:**
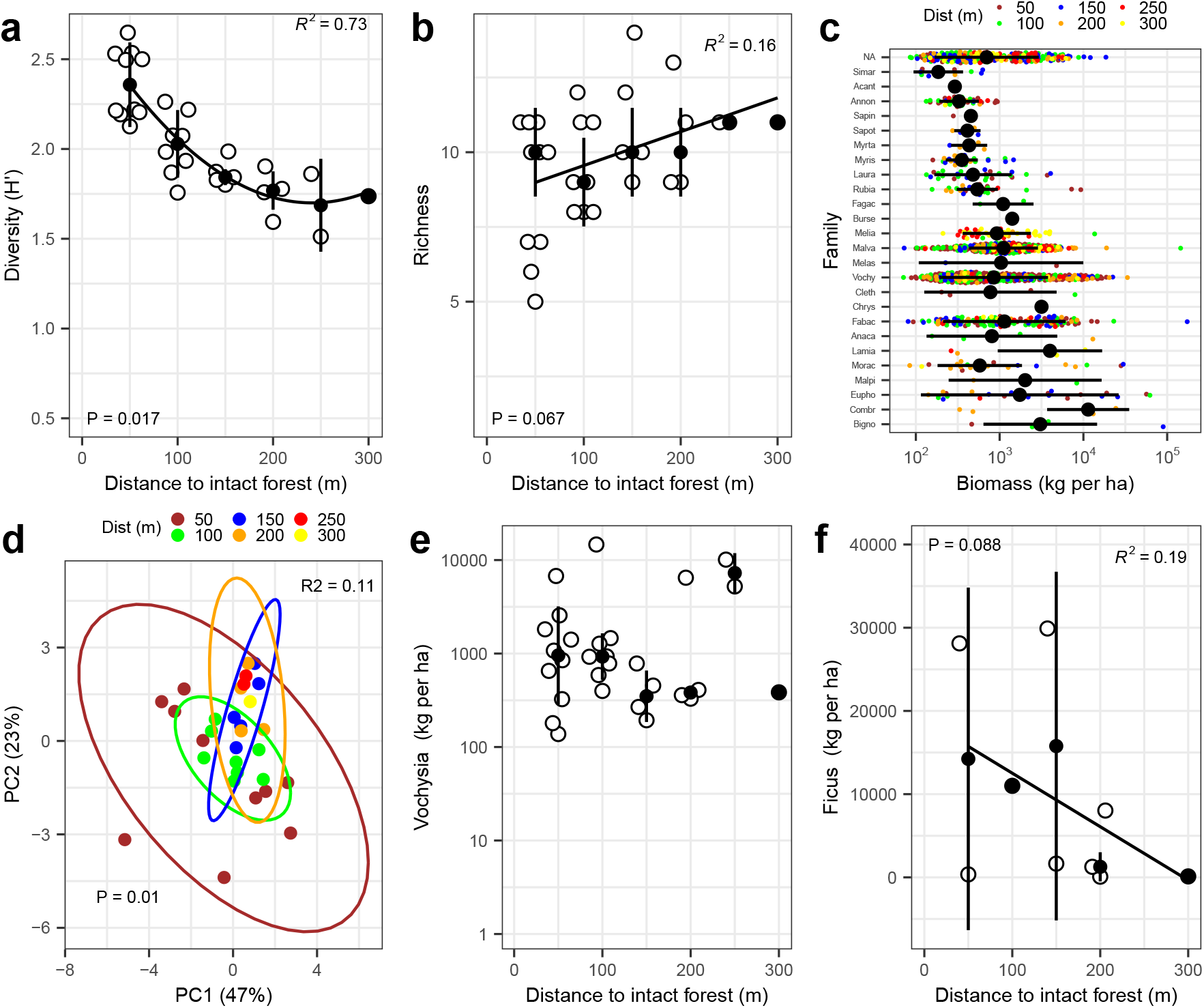
Taxonomic composition measures including (a) diversity, (b) richness, (c) overall biomass, (d) beta diversity, and (e,f) biomass of key taxa. White and colored dots with border show plot medians, black dots show plot-level data binned into discrete distance strata medians ± 1 absolute deviation, and black lines show linear or non-linear regression through strata medians.

### Traits

Overall trait regeneration highlighted successional stage associations over primary dispersal mode along distance to primary forest (Fig 4). Taxa associated with both early and late successional stages decreased significantly (p=0.042) and curvi-linearly by ~ −584.87 ± 167.31 kg per m with distance to edge explaining *~14*% variance among distance strata medians (Fig 4a). Dispersal modes did not show consistent trends in biomass with increasing distance to edge (Fig 4b).

~~~
## |                                                                                         |
## |                                                                                         |
## |                                                                                         |
~~~

**Figure 4:**
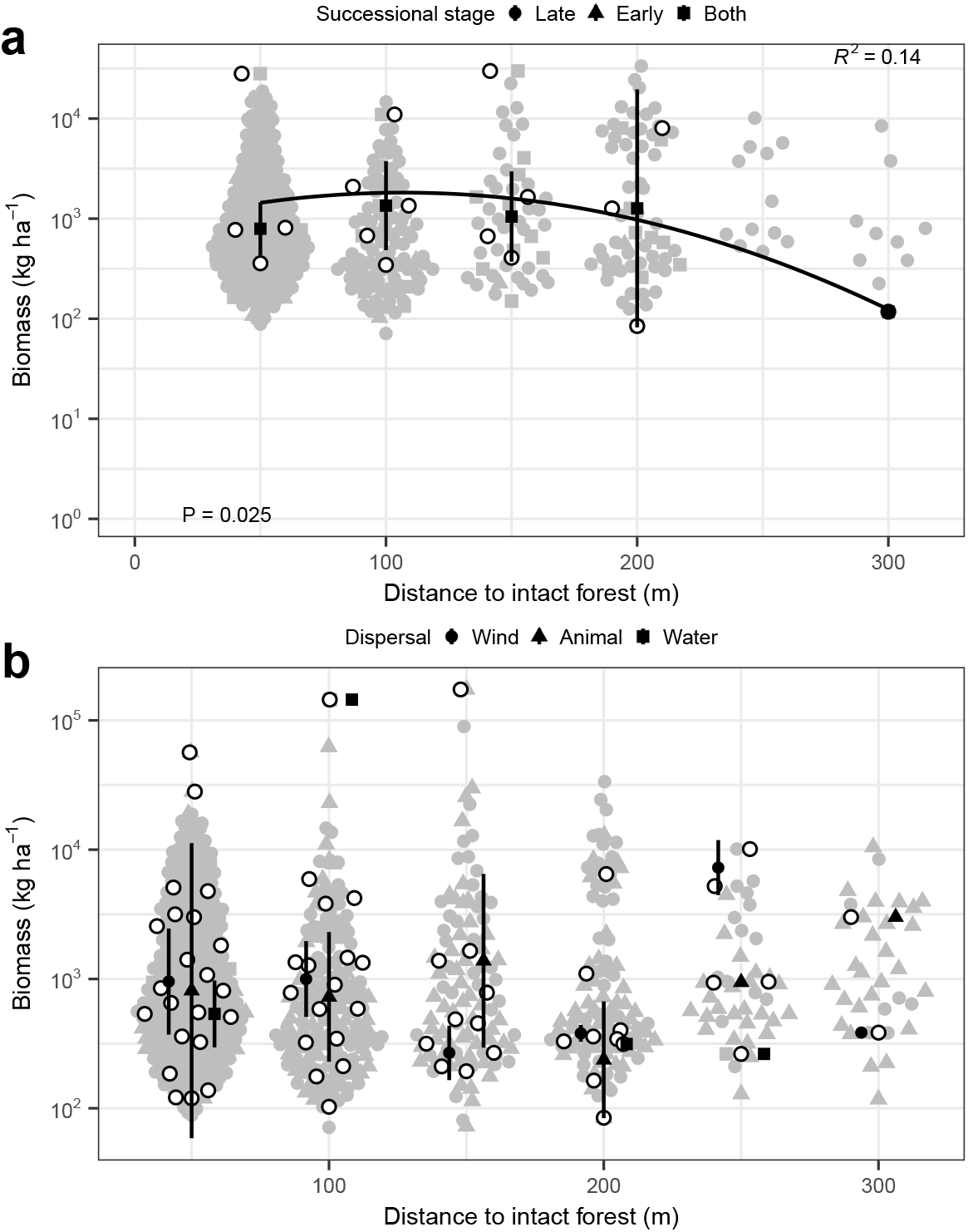
Functional regeneration based on taxa associations with (a) successional stages and (b) primary dispersal mode. Grey dots show individual tree values, white dots with border show plot medians, black dots show plot-level data binned into discrete distance strata medians ± 1 absolute deviation, and black lines show linear or non-linear regression through strata medians.

## Discussion

This study found significant edge effects on secondary forest stand wood and stem density, tree diversity and community composition, and some functional traits, yet not on overall aboveground biomass. It was initially hypothesized that community composition would vary with distance to high-quality intact primary forest edge, favoring more shade-tolerant taxa, due to light availability and dispersal potential. The evidence presented here supported edge effects on community composition and biomass of taxa that were associated with both early- and late-successional stages, but without detectable roles for light availability or general dispersal mode. Accordingly, this suggests that edge effects do significantly change humid tropical forest community taxonomic and functional composition, but primarily due to factors other than light competition or dispersal limitation (*Krishnadas et al., 2020*), and also suggests that aboveground biomass and even low taxonomic richness are resilient after a decade of regeneration.

Aboveground biomass storage is a key global ecosystem function and service, but edge effects tend to release about one-third of carbon lost from tropical deforestation (*Brinck et al., 2017*), with more expected in the future (*Mitchard, 2018*). Management has been found to explain most of biomass variation among tropical forest fragments, and wood density ~10% biomass variation (*Pyles et al., 2022*). Additionally, this study contributes that management via edge effects can also mediate ~10% of wood density variation without affecting local biomass, although median stem wood density can describe regional spatial biomass patterns (*Baker et al., 2004*). In this study, wood density and richness values may have been pulled down by early-successional clonal trees closest to the primary forest, which tend to have lower wood densities, lowering carbon (*Gonzalez J and Fisher, 1998; Pyles et al., 2022*), and increasing nearby stem density. Lower wood density can also reflect more dynamic forest stands (*Malhi et al., 2006*) with higher tree turnover, a consequence of forest fragmentation (*Nascimento and Laurance, 2004*) together with other global change drivers (*Laurance et al., 2014*), and may also reflect possible differences in soil fertility (*Malhi et al., 2004*), all of which represent possible future research directions, beyond strictly neutral dynamics (*Terborgh et al., 1995*). Furthermore, light availability at the plot level did not affect biomass storage, but instead, individual light gaps may be more important for biomass dynamics (*Chazdon and Fetcher,1984*).

Edge effects have been shown in other studies not only to lower forest biomass, but also to change community composition (*Anderson et al., 2022*), and in addition this study contributes that the decline in diversity can be rapid and non-linear across edge gradients, along with supporting the likelihood of changes to community composition lasting decades. This rapid loss of diversity across the edge gradient may be in part due to existing higher yet constant levels of shade limiting shade-intolerant seedling recruitment and/or performance, together with insufficient time for slower-growing shade-tolerant taxa to accumulate significant amounts of biomass. Another possible factor may be Janzen-Connell processes (*Terborgh, 2020; Wills,2006*), including conspecific negative density-dependence of seedling survival on basal area (*Comita and Hubbell, 2009*), as well as overall short dispersal kernels, whose effects can be mediated by shade (*Comita et al., 2014*) and in part by higher fungal pathogen pressure near conspecifics (*Jia et al., 2020*). While both diversity and community composition changed significantly with distance to intact forest in this study, the biomass of most individual taxa, surprisingly, tended to stay the same along the edge gradient. This lack of significant individual taxon biomass responses along this edge gradient could be due to stochastic population factors affecting their biomass growth, perhaps widening variability. Regardless of the underlying process, these absent patterns do also help explain the lack of pattern observed with plot biomass overall across the edge gradient. This explanation points to future studies of taxon-specific responses to edge effects, to help explain overall forest biomass dynamics near habitat edge. More specifically here, the biomass of the most abundant taxon *Vochysia* was resilient to edge effects in this study, however, that of the taxon *Ficus*, key for frugivores like birds and bats (*Cottee-Jones et al., 2015; Rafidison et al., 2020*), was significantly lowered across the secondary forest edge, pointing to potentially different responses of biomass and local food web associated ecosystem services. Results suggest that this pattern is specific to this locally widespread *Ficus* genus, since the overall biomass of other animal-dispersed taxa as a whole tended to stay the same across the edge gradient studied here. Interestingly, community composition also appeared to show lower variance across the edge gradient, although testing this observation in a forested area with evenly distributed area among distance strata bands would provide additional support.

Characterizing community composition by functional traits is also increasingly studied (*Kearney et al., 2021*), but less so regarding edge effects. A study in Madagascar found lower phylogenetic diversity closer to degraded forest edge, but no change in aboveground biomass (*Razafindratsima et al., 2018*). This study presents similar results in showing lower taxonomic diversity, although biomass specifically of taxa associated with both early- and late-successional stages also decreased with away from primary forest edge. This resulting decrease by successional stage could be explained by slightly lower habitat quality inferred near exposed habitat edges near roads, and accordingly relatively higher habitat quality near closed primary forest edge (*Ries et al., 2004*). While lower biomass of succession-agnostic taxa further from primary forest edge appears consistent with the expected forms of group-level dispersal kernels as an explanation, the lack of significant pattern along forest edge when binned by broad dispersal mode that was found here may limit the potential of seed dispersal to explain biomass trends. Instead, seedling survival may have been more important than dispersal in accumulating basal area and biomass (*Comita and Hubbell, 2009*). For taxa primarily dispersed by animals instead of wind, this could be explained by potentially broader local limitations in animal dispersal activity or abundance. Additionally, the potential for nearby dispersal to facilitate reforestation could be tested in future studies by measuring taxon-specific dispersal kernels.

Overall, this study highlights how abandoned wet tropical timber plantation can regenerate in alongside fragmented forests. Results support related syntheses that tropical forest biomass and taxonomic richness often regrows relatively quickly (*Davies et al., 2021*), while taxonomic composition recovers much more slowly, if not diverging altogether (*Norden et al, 2015*). These findings can suggest that more efficient reforestation could more actively target biodiversity conservation over other resilient functions like carbon storage, and that restoration efforts could include focusing activities like native species planting efforts on degraded habitat edges away from primary forest edges, and investing less in areas closer to existing forest edges, where recruitment may already maintain diversity levels. As forests continue to become fragmented (*Haddad et al., 2015*), understanding how to work with natural regeneration patterns around edge effects will likely become increasingly important.

## Acknowledgements

Thanks to Brandeis University Professor Dan Perlman for initial networking; Lehigh University Professor Don Morris and student Erin Lau for field support; concurrent Osa Conservation general manager Manuel Ramírez and Piro Biological Station staff Juan Carlos Cruz Díaz, Annia Barrantes, Larry Villalobos, Hansel Vargas, and visitors for field support; James Fifer, Chau Ho, and Michelle Spicer for review of early draft content; and Luisa Valdez for translation assistance.

Authors declare no conflicts of interest.

## Author contributions

NM, MV and AJ contributed to study conceptualization, design, administration, and supervision; NM and AJ contributed to funding acquisition; NM, AM, and EC, contributed to key data collection; NM and MV contributed to data analysis; NM wrote initial draft and later revisions.

Study team includes and recognizes diverse and historically-excluded contributions to research; with bilingual communication efforts completed and pending.

## Data statement

Code stored at github.com/nmedina17/osa (*Medina, 2022b*) with data stored at https://doi.org/10.5281/zenodo.7406478 (*Medina, 2022c*).

